# Assessing whole-cortex excitability from electromagnetic brain signals

**DOI:** 10.64898/2026.01.31.703018

**Authors:** Giovanni Pellegrino, Gian Marco Duma, Anna-Lisa Schuler, Meaghan Daub, Daniele Marinazzo, Giorgio Arcara, Andrea Soddu, Javier Rasero

## Abstract

Cortical excitability, the propensity of neural circuits to respond to internal or external perturbations, is a fundamental property of brain functioning, shaped by local microcircuitry, large-scale networks, and neurochemical architecture. In humans, excitability is inferred from multiple indirect metrics derived from spontaneous and stimulus-driven electromagnetic activity, yet it remains unclear whether these measures reflect a common underlying construct or distinct physiological processes. Here, we systematically compared ten previously validated excitability metrics using whole-head magnetoencephalography (MEG) recorded at rest and during 40 Hz auditory stimulation in a large sample of healthy adults. The measures spanned stimulus-driven synchronization, spectral power, *1/f* activity (hereafter defined as aperiodic), signal complexity, long-range temporal correlations, and phase–gamma synchronization. Hierarchical clustering revealed six separable excitability dimensions with limited redundancy, demonstrating that cortical excitability is inherently multidimensional. Aperiodic and alpha-band measures showed the strongest mutual coupling and the furthest spatial correspondence to stimulus-related responses, whereas temporal, complexity, and synchronization-based metrics were largely independent. The similarity between measures varied across cortical regions and functional networks, with maximal cluster separability in integrative and internally driven regions. These regions included orbitofrontal, temporal, anterior cingulate, and default mode areas, as well as visual and somatomotor networks. Finally, excitability dimensions showed distinct relationships with cortical morphology and neurotransmitter receptor density, implicating heterogeneous neurobiological substrates. Together, these findings provide a unified framework for interpreting excitability metrics and highlight the need for multimodal approaches when probing cortical excitability in health and disease.

## INTRODUCTION

Excitability refers to the propensity of neurons and neural circuits to respond to internal or external perturbations (*1*, *2*). Maintaining a balanced level of excitability is essential for stable cortical function, enabling reliable sensory processing and efficient information transmission (*3–5*). Dysregulation of excitability can disrupt this balance, impacting electrical, hemodynamic, and metabolic brain processes (*6–8*). Excitability is shaped by biological constraints acting at both local (region-specific) and global (whole-brain network) levels. These constraints encompass cortical morphology, genetic and neurochemical organization (*9–12*). Altered excitability balance has been implicated in the pathophysiology of numerous neuropsychiatric disorders, including stroke, epilepsy, amyotrophic lateral sclerosis, Alzheimer’s disease, and schizophrenia (*13–17*) . Accordingly, the ability to reliably measure excitability is critical for understanding normal and pathological brain function and has direct relevance for clinical care (*18*).

In humans, excitability has classically been assessed through neuronal responses to controlled perturbations, with non-invasive transcranial magnetic stimulation (TMS) providing a benchmark when assessing motor systems (*2*, *19*). Despite its utility, TMS is inherently limited in its capacity to probe excitability across distributed, whole-brain cortical networks (*18*). Electromagnetic responses evoked by 40 Hz auditory stimulation represent an alternative approach to assessing excitability. The 40 Hz auditory stimulation activates widespread cortical areas, with a right-lateralized temporo-central emphasis, and offer a stimulus-driven measure of excitability that likely reflects parvalbumin-positive interneuron activity and GABAergic–glutamatergic transmission(*20–23*). Beyond stimulus-driven paradigms, considerable effort has been devoted to developing stimulus-independent approaches for mapping excitability across the whole-brain level (often referred to as intrinsic excitability) (*24–26*). These techniques leverage spontaneous electromagnetic activity such as parametrizing *1/f* distribution (hereafter defined as aperiodic) of power spectrum density, signal entropy, and gamma-band phase synchronization. These approaches are particularly attractive because they can be applied across the entire cortex without external stimulation, and because fluctuations in spontaneous brain activity are closely linked to changes in cortical excitability (*27*, *28*).

Despite their growing application, stimulus-independent measures of excitability have largely been studied in isolation and are rarely validated against one another, limiting the understanding of underlying mechanisms and clinical applicability. For example, in a previous study, classical TMS-based measures of motor cortex excitability correlated with only one of the many available intrinsic metrics (*24*). More broadly, little is known about the reliability, regional specificity, and complementarity of excitability measures, or about their relationship to cortical architecture, genetic factors, and neurochemical organization (*18*). Together, these observations suggest that commonly used excitability metrics may differ in their physiological specificity, regional expression, and underlying biological constraints. To address this gap, we systematically examined stimulation-based and intrinsic excitability measures in a large sample of healthy individuals using whole-head MEG data acquired both at rest and during 40 Hz auditory stimulation.

We hypothesized that: (1) different excitability measures exhibit varying degrees of association; (2) their correspondence depends on cortical region and network organization; and (3) rather than reflecting a single ground truth, distinct measures capture complementary aspects of cortical excitability and show differential relationships with cortical structure and receptor density. By integrating functional, structural, and neurochemical dimensions, this work provides a biologically grounded framework for interpreting excitability measures and advancing their potential as biomarkers of brain function and dysfunction.

## RESULTS

### Divergent excitability estimates from identical brain signals

MEG data were acquired from 53 participants using a 275-channel whole-head system. Each participant completed two six-minute eyes-closed recordings (rest and 40 Hz auditory stimulation) in randomized order. After source reconstruction, we extracted ten excitability metrics from 100 cortical regions (Schaefer atlas (*29*)): eight from resting-state activity and two from the 40 Hz stimulation condition. These metrics included the Excitability Index (corresponding to the spatial phase synchrony in gamma band) (*30*), spectral aperiodic exponent and offset (*31*), sample entropy (*32*), detrended fluctuation analysis (DFA) (*33*), three alpha-band measures (absolute, relative, and oscillatory power (*34*)), and inter-trial phase consistency (ITPC) during 40 Hz stimulation and baseline-corrected ITPC (*35*). The Fig.1 depicts the group-averaged spatial distribution of these excitability measures. To note, metrics were directionally adjusted so that higher values uniformly reflect higher excitability (see Methods).

**Figure 1.**
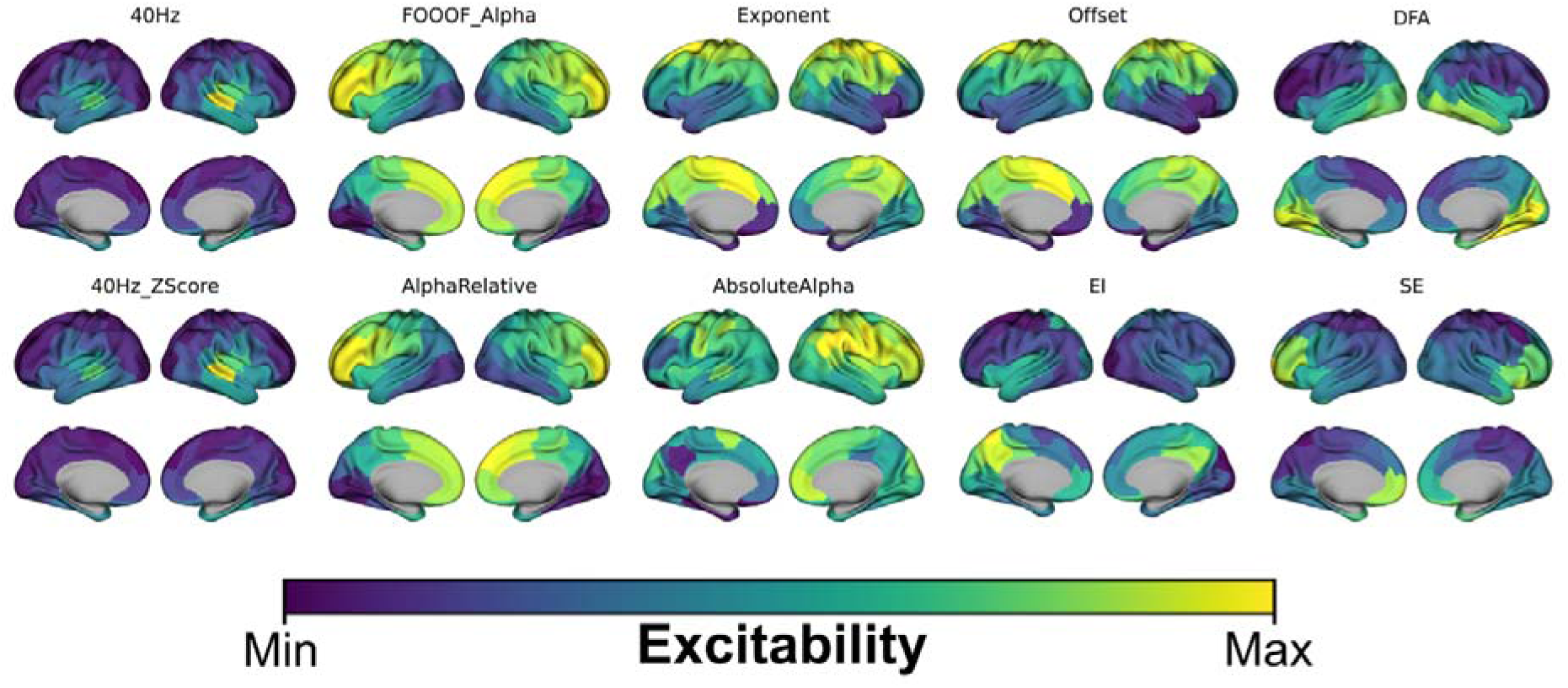
Spatial distribution of cortical excitability measures. Group-averaged cortical maps showing the spatial distribution of each excitability measure across the brain. Measures are displayed separately, with color indicating the magnitude of the corresponding metric, averaged across participants and mapped onto the cortical surface. DFA ≡ detrended fluctuation analysis; EI ≡ Excitability Index; SE ≡ Sample Entropy.

### Global relationships among excitability measures

To assess relationships among metrics, we computed Spearman correlations between all pairwise combinations of the ten whole-brain excitability measures, yielding a 10 × 10 similarity matrix per scan. Averaging across scans and converting similarity to distance (D = 1 − ⟨R⟩), hierarchical clustering revealed six distinct clusters (Fig. 2a). Cluster number was selected based on the largest minimum inter-cluster distance (d = 1.05) and supported by a silhouette score of 0.60, indicating well-defined groupings. Each cluster comprised measures derived from different signal properties and analytical approaches, indicating limited redundancy among metrics and suggesting that they capture complementary aspects of excitability rather than a single underlying construct. ***Cluster I*** included stimulus-driven synchronization measures (ITPC 40 Hz and baseline-corrected ITPC), ***Cluster II*** captured long-range temporal correlations measured from detrended fluctuation analysis (DFA), ***Cluster III*** corresponded to the Excitability Index, ***Cluster IV*** comprised sample entropy, ***Cluster V*** included aperiodic spectral features (exponent and offset), and ***Cluster VI*** contained alpha-band measures (absolute, relative, and oscillatory power) (see Fig1A). The strongest inter-cluster spatial similarity was observed between alpha-based and aperiodic clusters (Fig. 2b), consistent with their shared spectral origin. In contrast, DFA, entropy, and stimulus-locked measures showed weak correspondence with spectral metrics. Finally, the stimulus-driven cluster exhibited small-to-moderate and non-significant (all spin-permuted *p* > .05) spatial similarity values with the rest of clusters, including both positive (SE: ρ = .359, DFA: ρ = .328, EI: ρ = .148) and negative (Alpha: ρ = -.153) associations. The strongest association (in magnitude) was found with the aperiodic cluster (ρ = -.591, spin-permuted *p* = .021), yet this effect did not survive False Discovery Rate (FDR) correction (i.e. *padj* > .05). These results demonstrate that excitability measures derived from identical neural signals diverge substantially, supporting Hypothesis 1 that different metrics exhibit variable association patterns.

**Figure 2.**
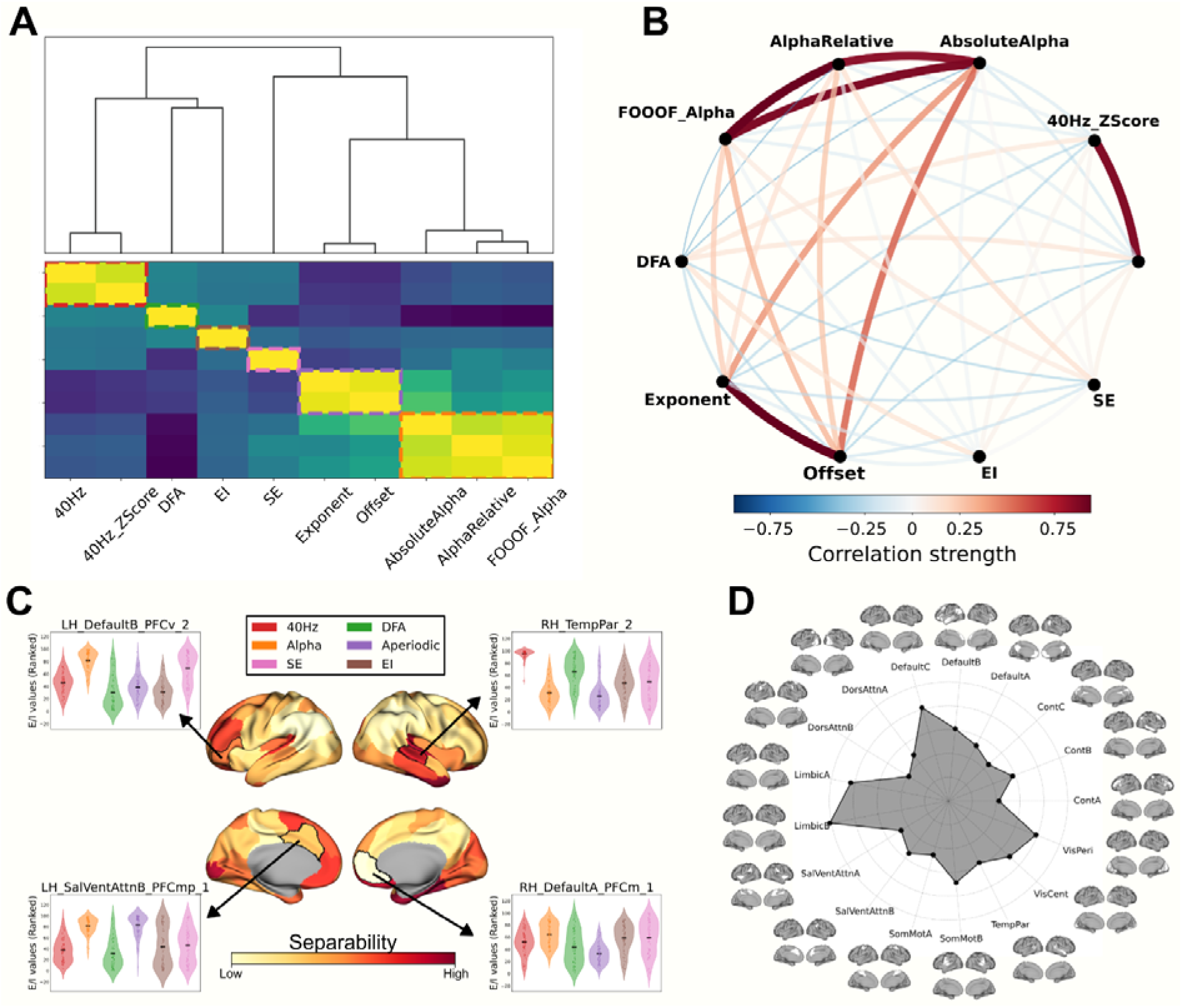
Clustering and separability of excitability measures. **(A)** Hierarchical clustering of the ten excitability measures revealed six distinct clusters: stimulus-driven oscillatory responses (Cluster I), detrended fluctuation analysis (Cluster II), a gamma-band phase synchrony (Excitability Index) (Cluster III), signal complexity (Sample Entropy) (Cluster IV), aperiodic spectral features (Cluster V), and alpha-band measures (Cluster VI). **(B)** spatial similarity matrix showing stronger inter-cluster relationships between alpha-based and aperiodic measures than between temporal/complexity measures and spectral measures. **(C)** Regional separability (F-values) indicating cortical areas that contribute most to cluster differentiation. **(D)** Network-level separability shown for standard resting-state networks mapped onto the Schaefer atlas.

### Regional and network specificity of excitability cluster coupling

We next examined whether cluster structure varied across cortical regions and functional networks. Cluster-specific centroid maps were generated by averaging ranked excitability maps across measures within each cluster. One-way ANOVA F-values quantified separability across clusters within each cortical region and network, with higher F-values indicating stronger differentiation (see Methods). Cluster separability was strongest in orbitofrontal cortex, anterior cingulate, visual cortex, cuneus, precuneus, and temporal poles (Fig. 2c). Early visual regions showed pronounced divergence between spectral, stimulus-locked, and broadband measures, while the precuneus exhibited strong separability driven by long-range temporal correlations, complexity, and aperiodic activity. Network-level analyses showed that separability was most pronounced in limbic and default-mode networks, consistent with their variable and context-dependent dynamics, followed by visual and somatomotor networks (see Fig. 2D) (*36*). These findings indicate that excitability metrics vary across cortical systems (*10*), supporting Hypothesis 2 that correspondence among measures depends on cortical region and network organization.

### Local vs. distributed contributions to cluster coupling

To determine whether cluster relationships reflect local effects or distributed patterns across the cortex, we modeled pairwise cluster associations within each brain parcel using linear mixed models with subject as a random effect and FDR correction for multiple comparisons. Local cluster–cluster analyses focused on positive correlations, which reflect spatial co-expression of excitability dimensions and identify regions where multiple measures are jointly modulated. Such positive associations are most consistent with shared underlying physiological mechanisms. Statistical inference was nonetheless based on two-tailed tests, providing a conservative assessment of significance while restricting interpretation to physiologically meaningful positive effects. We report z-statistics reflecting the strength of associations between clusters (Fig. 3). A near whole-brain positive correlation was observed between alpha-based and aperiodic clusters (*padj* < .05; Fig. 3), indicating that alpha oscillations co-vary with scale-free dynamics at a global level. Strong local correlations were particularly evident in temporal and sensorimotor regions. The alpha cluster also showed a more diffuse correlation with entropy, with the strongest effects in dorsolateral prefrontal cortex, temporo-parietal junction, and motor cortex. In contrast, associations among the remaining cluster pairs were spatially sparse or negligible across cortical parcels. The absence of widespread regional correspondence suggests that most excitability clusters are largely independent at the local level, supporting a multidimensional organization of excitability in which distinct metrics capture complementary physiological processes operating across different spatial and temporal scales.

**Figure 3.**
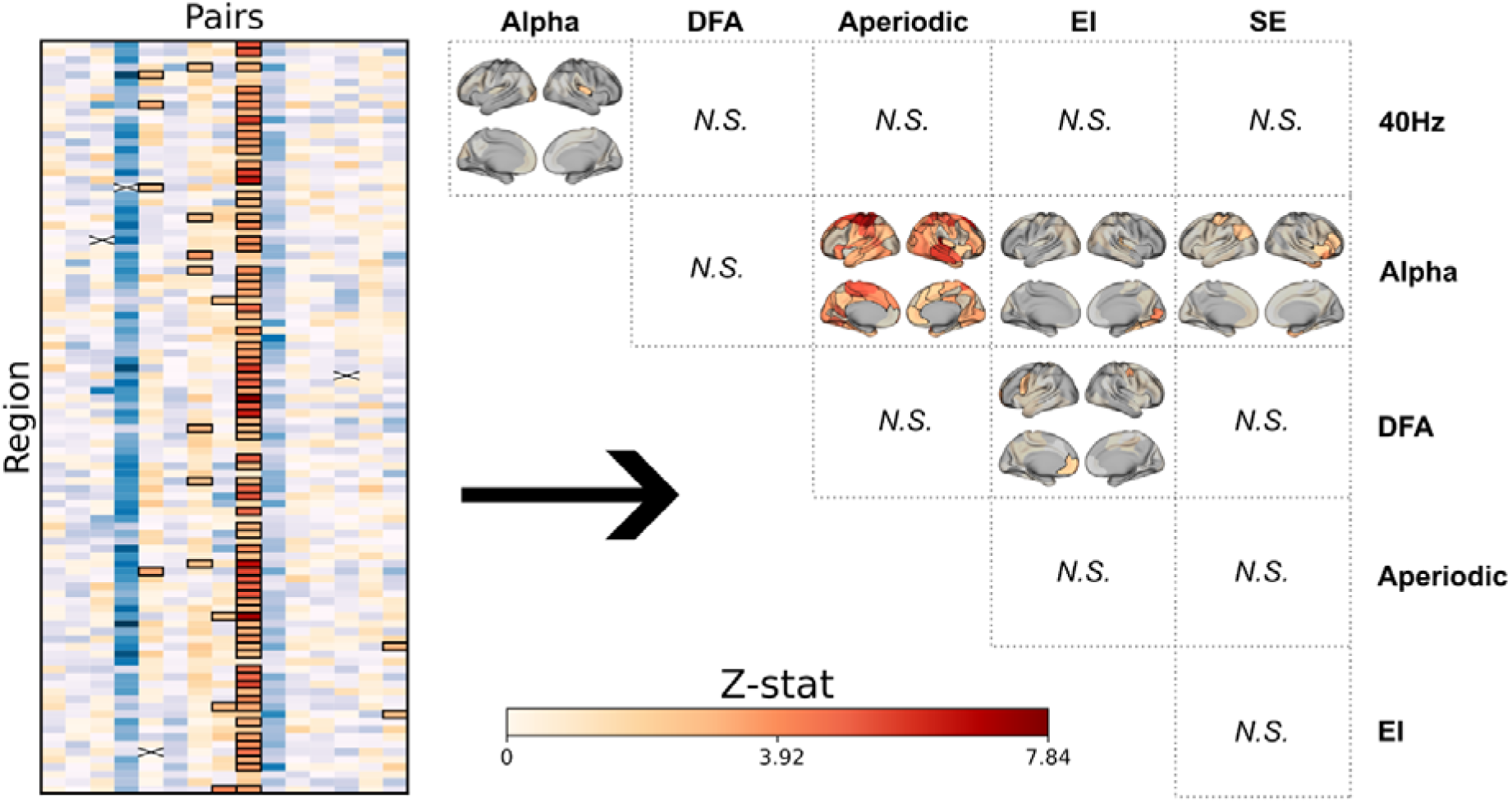
Spatial organization of positive relationships between excitability measures. **Left**, region-by-measure heatmaps showing pairwise associations between excitability clusters, expressed as z-statistics from linear mixed-effects models accounting for repeated measures; significant positive associations are highlighted in squares. **Right**, cortical maps showing the spatial distribution of statistically significant positive correlations between cluster pairs (padj < 0.05). A near whole-brain coupling is observed between alpha-based and aperiodic excitability clusters, with particularly strong local correlations in temporal and sensorimotor cortices. More spatially circumscribed correlations are evident between alpha-based and entropy-based clusters, most prominently in dorsolateral prefrontal, temporo-parietal junction, and motor regions. Other cluster pairings show weak or spatially isolated correspondence, indicating largely independent local excitability mechanisms across cortical parcels. DFA detrended fluctuation analysis; EI Excitability Index; SE Sample Entropy.

### Excitability measures show differential associations with cortical structure

Brain structural and neurochemical architecture are known to have a direct influence on cortical excitability. To this end, we investigated the relationship between cortical thickness, receptor density maps and the extracted excitability clusters, assessing their association as global and local structural correlates of intrinsic excitability. At the global (whole-brain) level, for each scan, we computed Spearman correlations between cortical thickness and each cluster map. To test whether the association of each cluster with thickness deviated significantly from zero across scans, we compared the mean observed correlation to a null distribution generated by averaging scan-wise spin-permuted correlations, thereby accounting for spatial autocorrelation. Although none of these Associations survived FDR correction (all *p* > .05), we observed a positive average spatial correlation of thickness with 40 Hz (ρ = .291), alpha-based (ρ = .229) and sample entropy (ρ = .093) clusters, and a negative correlation with the aperiodic (ρ = - .243), DFA (ρ = - .111), and EI (ρ = - .026) clusters. These trends indicate heterogeneity in both the magnitude and direction of association between excitability measures and cortical thickness.

At the local level, we assessed the association between cortical thickness and excitability values for each cluster within each region using separate linear mixed models, followed by FDR correction for multiple comparisons. As shown in Figure 4A, the 40 Hz cluster exhibited a significant positive association with thickness in the right ventral prefrontal cortex. For the alpha-based cluster, a positive association was found in the right orbitofrontal cortex, alongside a negative association in the left lateral prefrontal cortex. For DFA, significant associations were identified in the right extrastriate visual (positive) and right posterior cortices. The aperiodic cluster showed only negative associations, localized in the left precuneus and the right dorsal prefrontal cortex. Notably, despite exhibiting the weakest global correlation with thickness, the Excitability Index showed the largest number of significant local associations, including positive effects in the right medial prefrontal cortex and negative effects in the left ventral prefrontal cortex and temporal pole. Sample Entropy showed significant positive associations in the right medial prefrontal cortex and the left extrastriate visual cortex. Together, these results show heterogeneous structural correlates across clusters, supporting Hypothesis 3.

**Figure 4.**
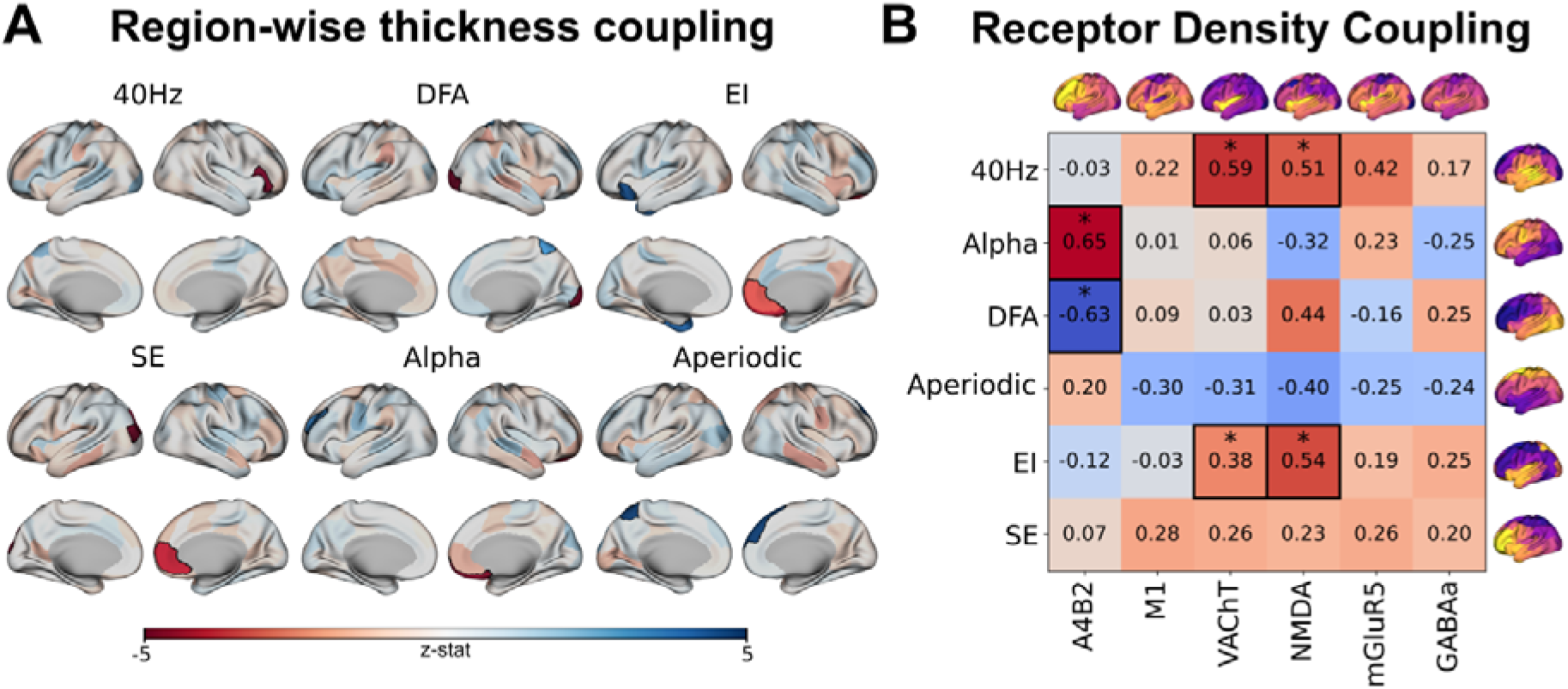
Structural and neurochemical correlates of excitability clusters. **(A)** Patterns of region-wise associations between cortical thickness and each excitability cluster. The highlighted regions with the black border represent the brain regions showing a cluster-thickness correlation after FDR correction. We report z-statistics derived from the linear mixed model. **(B)** Heatmap showing spatial correlations between excitability clusters (averaged across subjects) and cortical receptor density maps, illustrating the neurochemical profiles associated with each excitability dimension. DFA detrended fluctuation analysis; EI Excitability Index; SE Sample Entropy.

### Excitability clusters reflect cortical neurochemical architecture

To examine neurochemical correlates, we computed spatial Spearman correlations between cluster maps and PET-derived receptor density maps, focusing on GABAergic, glutamatergic, and cholinergic systems (Fig. 4B). Significance was assessed using spin permutation tests, and multiple comparisons corrected for by FDR. Glutamatergic and cholinergic markers showed the strongest associations. Specifically, NMDA receptor density correlated positively with the Excitability Index (ρ = .538, *padj* = .004) and 40 Hz responses (ρ = .505, *padj* = .016). VAChT density also correlated positively with the Excitability Index (ρ = .381, *padj* = .046) and 40 Hz cluster (ρ = .589, *padj* = .004). The alpha-based cluster correlated positively with α4β2 receptor density (ρ = .655, *padj* = .012), while the DFA cluster showed a negative correlation with α4β2 density (ρ = - .634, *padj* = .045). These findings indicate that excitability clusters are differentially embedded within neurochemical architecture, with stimulus-locked and rhythmic excitability linked to glutamatergic and cholinergic systems and scale-free dynamics showing an opposing profile, further supporting Hypothesis 3.

## DISCUSSION

In this study, we systematically compared a broad set of MEG-derived excitability metrics, including both stimulus-dependent and stimulus-independent measures. Using a unified framework applied to resting-state activity and 40 Hz auditory responses, we assessed the extent to which these proxies converge, how their relationships vary across regions and functional networks, and whether they exhibit distinct structural and neurochemical signatures. Our findings support a multidimensional view of cortical excitability, in which different metrics capture complementary aspects of neural dynamics rather than reflecting a single underlying construct. Although each metric has been individually validated in prior work (*16*, *25*, *30*, *33*, *34*, *37*), their cross-validation and shared physiological interpretation have remained uninvestigated. In this study, we addressed this gap by directly comparing multiple excitability metrics within the same dataset and identifying their differential relationships to cortical structure and neurochemical organization (*38*). To our best knowledge, this is the first study directly comparing multiple excitability metrics within the same dataset and identifying their differential relationships to cortical structure and neurochemical organization.

### Excitability as a Multidimensional Construct

Hierarchical clustering of the ten excitability metrics revealed six distinct clusters (Fig. 2A), supporting Hypothesis 1 that different metrics exhibit variable association patterns. Specifically, Cluster I grouped stimulus-driven synchronization measures (ITPC 40 Hz and baseline-corrected ITPC), Cluster II captured long-range temporal correlations (DFA), Cluster III corresponded to the Excitability Index, Cluster IV comprised sample entropy, Cluster V included aperiodic PSD features (exponent and offset), and Cluster VI contained alpha-band measures (absolute, relative, and oscillatory power). The strongest inter-cluster similarity was observed between alpha-based and aperiodic clusters, whereas DFA, entropy, and stimulus-locked measures showed weak correspondence with spectral metrics (see Fig. 2B). The cluster separation between stimulus-locked and intrinsic measures is central for interpreting excitability metrics. Rather than simply reflecting stimulus-driven responsiveness, intrinsic measures appear to index internal regulatory dynamics that shape the baseline cortical state. Within intrinsic metrics, alpha-based measures clustered tightly and were closely associated with aperiodic spectral features, whereas entropy and DFA remained more independent. Given that these metrics derive from different signal features, they are likely to reflect partially independent physiological mechanisms operating across spatial and temporal scales, rather than providing redundant estimates of a single construct. These findings support a multidimensional framework in which different metrics probe complementary aspects of cortical dynamics.

### Global and Local Specificity of Excitability Relationships

Regional and network patterns of separability indicate that differences between excitability dimensions are amplified in areas with distinct functional roles and intrinsic dynamics. Integrative and internally driven regions, including the orbitofrontal cortex, anterior cingulate, precuneus, and temporal poles showed strong separability, consistent with their high variability and flexible modulation of neural activity (*39*, *40*). By contrast, in posterior sensory areas like the visual cortex, different excitability measures often disagreed with one another. This likely reflects the precise and specialized organization of the sensory cortex, where tightly controlled oscillations and layered circuitry cause each measure to capture a different aspect of neural responsiveness (*41–43*). Notably, the precuneus emerged as a major contributor to separability, likely due to its central role within the default-mode network, where excitability is predominantly shaped by intrinsic, large-scale processes rather than immediate sensory input, allowing multiple excitability dimensions to dissociate (*44*, *45*) . At the network level, pronounced separability observed in limbic, default-mode, visual, and somatomotor systems further supports this interpretation. Together, these findings suggest that excitability dimensions are differentially expressed across networks operating at distinct temporal scales and reflect both local circuit properties and large-scale dynamics, rather than a single uniform process.

At the parcel level, relationships between excitability clusters were largely selective, with a notable exception being the coupling between alpha and aperiodic clusters widespread across the brain. This likely reflects their shared sensitivity to excitation–inhibition balance, with alpha oscillations more involved into inhibitory control and gating, and aperiodic activity indexing broadband synaptic dynamics (*46–48*). The strongest associations were observed in temporal and sensorimotor cortices, consistent with dense recurrent connectivity and organized inhibitory networks in these regions (*49–51*). Alpha measures also showed positive coupling with entropy, particularly in dorsolateral prefrontal, temporoparietal, and motor regions, linking rhythmic activity to neural complexity in executive, attentional, and motor related brain regions(*52–54*). In contrast, most other cluster pairings showed weak or localized correspondence, supporting Hypothesis 2 that inter-metric relationships depend on cortical region and functional system.

### Distinct Structural and Neurochemical Signatures

A key contribution of this work is the integration of excitability metrics with cortical structure and neurochemical architecture, directly addressing Hypothesis 3. Parcel-wise analyses indicate that the relationship between cortical thickness and excitability is spatially heterogeneous, varying in both magnitude and direction across clusters. These patterns suggest that cortical thickness sustains different excitability dimensions through region-specific mechanisms, consistent with the idea that structural constraints interact with functional specialization (*11*, *12*). The association between thickness and excitability frequently showed opposing directions within the same cluster, implying that cortical thickness does not exert a monotonic influence on excitability. Instead, thickness may reflect how local architecture supports distinct physiological processes, depending on regional circuit organization(*55–57*). In this view, thickness could index structural capacity that facilitates certain excitability mechanisms in some cortical contexts while constraining or reshaping them in others (*58*, *59*) . Importantly, the observation that some clusters exhibit minimal global associations yet robust local effects highlights a critical dissociation between global structural correlates and local structure–function relationships.

Neurochemical associations further elucidate the physiological basis of excitability clusters. Stimulus-evoked and Excitability Index-based clusters correlated positively with NMDA and acetylcholine receptor densities, consistent with their roles in supporting synaptic gain, gamma-band synchronization, and heightened cortical responsiveness (*60*, *61*). The alpha cluster was associated with α4β2 nicotinic acetylcholine receptors, linking rhythmic inhibitory modulation to cholinergic signaling (*62*). Conversely, the DFA cluster showed a negative association with excitatory receptor density, supporting the idea that long-range, scale-free activity arises preferentially in regions with lower excitatory drive, likely reflecting emergent network-level properties. Overall, these findings support Hypothesis 3 by demonstrating that different excitability measures are grounded in distinct, and sometimes opposing structural and neurochemical dependencies, further emphasizing the need to consider spatial scale and functional context when relating brain properties to excitability.

### Implications for Excitability Measurement

The limited convergence among excitability metrics indicates that reliance on a single measure captures only a subset of excitability-related physiology. In practice, this suggests that studies using resting-state EEG/MEG should adopt a multimodal measurement strategy, combining at least one spectral metric (e.g., alpha power), one scale-free metric (e.g., aperiodic exponent or DFA), and one complexity measure (e.g., entropy). This approach minimizes the risk of over-interpreting any single excitability dimension and enhances robustness against methodological biases and metric-specific sensitivities (*24*).

### Limitations and Future Directions

Several limitations of the present study should be acknowledged. First, 40 Hz auditory stimulation provides a specific form of stimulus-driven excitation that may not generalize to all cortical regions or sensory modalities (*35*). Although TMS is the classical benchmark for excitability, its application outside the motor cortex can be unreliable, limiting its use as a universal reference (*18*). Second, MEG source reconstruction is biased toward superficial cortical sources, which may influence the observed spatial patterns (*63*). Third, receptor density maps provide only an indirect estimate of neurochemical architecture; density does not equate to receptor activation, and normative PET maps cannot capture individual variability. Finally, computational modeling integrating structure, function, and neurochemical data could help establish causal links between excitability metrics and underlying neuronal mechanisms, providing a more mechanistic interpretation of the observed relationships.

## Conclusion

In summary, our findings indicate that cortical excitability is a multidimensional construct, with different metrics capturing complementary aspects of neural dynamics. Although these measures diverge substantially, they exhibit systematic regional, structural, and neurochemical associations.

Together, these results provide a biologically grounded framework for interpreting excitability metrics and underscore the importance of multimodal integration for mapping cortical function and dysfunction.

## MATERIALS AND METHODS

### Participants and Experimental Design

This study complies with the Declaration of Helsinki and was approved by the Ethics Board of the Province of Venice. All participants signed a written informed consent before participation. We recruited 1) healthy adult subjects, 2) without hearing deficits, 3) without contraindication to MRI, 4) who did not take any neuropsychiatric medications, and 5) had no history of neuropsychiatric conditions. We included 53 individuals (40 female, mean age=29.5 ± 6.9) who underwent a total of 72 whole head MEG scans at rest and during stimulation with 40 Hz auditory sounds. All patients also had a brain anatomical MRI to characterize brain structure.

### MEG recording

MEG recordings were conducted in a shielded room using a 275-channel CTF system (MISL, Vancouver, Canada). Eye movements and EKG were recorded with bipolar electrodes, and data were sampled at 1200 Hz. Head position was continuously tracked via coils placed on anatomical landmarks (nasion and preauricular points). Participants maintained a regular sleep schedule for three days before the scan. The scan was performed with subjects lying supine, eyes closed, relaxed, and not attending to sounds. Resting-state and 40 Hz auditory stimulation (6 minutes each, randomized order) were recorded (*64*). Sounds were delivered binaurally via MEG-compatible tubes at 85 dB, using a 40 Hz amplitude-modulated 1000 Hz tone with 6 ms fade-ins/outs, generated in MATLAB. Sound levels were balanced, and no discomfort was reported. The 40 Hz sound was generated in MATLAB (The Mathworks) according to the formula

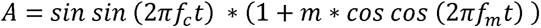

where *A* is the amplitude, *f_c_* is the carrier frequency set to 1000 Hz, *m* is the modulation depth, *f_m_* is the frequency of modulation, set to 40 Hz and *t* is the vector of time points for one sec of stimulus, at a sampling rate of 44100 Hz. Stimulation had 1-second duration followed by 1 second of silence, forming 2-second trials (*65*). A total of 180 trials were delivered over 6 minutes. Stimuli were delivered using PsychoPy (*66*). To ensure timing precision for gamma synchrony analysis, the actual sound was recorded via an MEG analog channel, allowing realignment of digital triggers (*67*).

### MEG preprocessing

MEG analysis was performed with the Brainstorm toolbox (*68*) operating with MATLAB (The Mathworks, v. 2016b). MEG data processing included the following steps: 1) third-order spatial gradient noise cancellation, 2) resampling to 600 Hz, 3) notch filter at 50Hz and harmonics (100, 150, 200 Hz), 4) highpass filter at 1Hz, 5) detection of cardiac activity from the EKG signal, followed by removal of EKG-related artifacts using Signal Space Projection (first component). For the 40 Hz auditory stimulation paradigm, data were segmented into 3-second epochs centered on stimulus onset (−1.5 to +1.5 seconds). Epochs were visually inspected, and those containing artifacts were rejected.

### Cortical source modelling

The brain MRI was performed for each participant with a 1.5 T Achieva Philips scanner (Philips Medical Systems, Best, The Netherlands). A 3-dimensional Magnetization Prepared Rapid Gradient Echo (MP-RAGE) T1-weighted scan was acquired using a 32-channel receiver head coil with the following parameters: repetition time [TR]=8.3 msec, echo time [TE]=4.1 msec, flip angle=8°, inversion time TI= 950 msec, isotropic spatial resolution=0.87 mm^33^. MRI segmentation was performed with the CAT12 toolbox (*69*) directly from Brainstorm. We obtained the following mesh surfaces: brain (15000 vertices), inner skull, outer skull and head. Sensor coregistration was performed on the head surface based on digitized headpoints acquired with the pholemus system. The forward model was computed using the overlapping spheres method. The inverse model was computed using weighted minimum norm estimate model (dipole orientation normal to surface).

### Excitability measures

All excitability measures were computed in the source space, at cortical level, for 100 parcels of the Yeo’s 17-network in the Schaefer parcellation (*29*) extracted for each subject with CAT12. The following measures were computed:

Stimulus-related metrics:

- *40Hz auditory stimulation:* We extracted Inter-Trial Phase Consistency (ITPC) between 39 and 41 Hz.. Higher ITPC are associated with higher gamma synchronization relative to stimulation (*21–23*, *65*). ITPC is bounded between 0 and 1. The higher the *ITPC* is, the higher is the excitability. *ITPC* was estimated for the entire time-course (−1.5 to +1.5 seconds) and for each vertex of the cortical mesh before averaging within parcels.
- *40Hz z-score* ITPC: ITPC values were converted into z-scores considering -500 to -200 ms as reference signal. This reference/baseline time-window was selected to avoid proximity to the sound and edge effects. ITPC z-transformed measures were averaged in the 300-700 ms time-window, as after about 200 ms of 40 Hz sounds brain activity reaches a steady cortical synchronization increase.

RS-Based metrics:

- Excitability Index: This index was introduced and validated by Meisel et al. (2015) (*30*). It corresponds to the mean spatial phase synchronization in specific frequency bands and it correlates with cortical excitability as estimated via perturbational approaches and modulated via antiseizure medications in epilepsy patients (*70*) . This estimate roughly corresponds to the well-known phase locking value (PLV), broadly exploited in neuroscience and neurology, computed across space rather than time. We selected the high gamma band (55-95Hz), as recommended in previous studies (*67*);
- *Exponent and offset of the aperiodic component of the Power Spectrum:* Converging evidence showed that the exponent of the power spectrum decay slope of electrophysiological signals, provides information on the E/I balance of the system (*25*, *31*). A steeper (higher or more negative) exponent indicates greater inhibitory activity, whereas a flatter (lower or less negative) exponent suggests increased excitation. To compute the exponent value we used the FOOOF implementation available in the Brainstorm toolbox (*71*) considering a frequency range between 1Hz and 40Hz, and extracted the exponent and offset of the 1/f slope as an excitability measure (*49*).
- *Sample Entropy:* Entropy measures have been associated with neural excitability, with lower entropy typically indicating higher excitability, and higher entropy reflecting reduced excitability (*32*, *72*). Sample entropy has proven to be particularly sensitive to changes in excitability (*37*). We computed sample entropy implementing Cannard & Delorme’s code (*73*) within Brainstorm.
- Detrended Fluctuation Analysis (DFA): DFA is a method used to measure long-range temporal correlations in time series data. In neuroscience, it helps assess the stability of brain activity by analyzing fluctuations in neural signals. Changes in DFA scaling can reflect shifts in excitability (higher DFA values = higher excitability), offering insights into both normal brain dynamics and neuropsychiatric/neurological disorders (*33*). The computation of DFA was performed in alpha band, following the method described in Van Nifterick et al. 2023 (*33*) adapting in Brainstorm the code available at https://github.com/annevannifterick/fEI_in_AD;
- *Alpha band power:* This measure is widely considered a marker of E/I, with higher alpha power corresponding to lower E/I (*34*). Alpha power (8-12 Hz) was computed with the Welch method, window length = 1 sec; overlap = 50%.

Crucially, to ensure that larger values consistently corresponded to higher excitability across the alpha-based, sample entropy and aperiodic metrics, their brain patterns, ***x***, were range-reversed (*x*→ *Max* + *Min* - *x*).

### Clustering and separability analysis

To assess similarity across the ten different excitability measures, we conducted a hierarchical clustering analysis. We computed Spearman’s correlation across all pairs of excitability measures, yielding a 10 × 10 similarity matrix (R) for each scan. Next, we calculated a distance matrix defined as **D = 1 −** ⟨**R**⟩, where ⟨R⟩ denotes the similarity matrix averaged across all scans. Hierarchical clustering was then performed on **D** using *SciPy* ’s linkage function with the average linkage method (*74*). To evaluate the contribution of individual regions to cluster separation, we computed a centroid map for each cluster by spatially averaging the ranked brain maps of the excitability measures included in that cluster. Ranked data were used to maintain consistency with the clustering analysis based on Spearman’s correlation. We then applied an ANOVA-based approach and, for each region, extracted the F-statistic from a one-way ANOVA. Larger F-values indicated greater differences in excitability measures across clusters for that region. These F-values were also averaged across regions within each of the 17 resting-state networks (RSNs), to facilitate interpretation of cluster separability from a functional organization perspective.

### Neurotransmitters Receptor density

We further examined the functional properties of each cluster by evaluating their association with neurotransmitters primarily involved in the regulation of the excitability. Whole-brain receptor density maps for neurotransmitters were extracted following the seminal work of Hansen and colleagues (*38*), which provided openly available volumetric neurotransmitter density maps derived from positron emission tomography (PET) using different ligands across a cohort of more than 1000 subjects. We then parcellated these receptor density maps to the Schaefer 17-network solution atlas of 100 brain regions. We considered the receptor density maps of the following neurotransmitters, which are known to be involved in regulation of excitability: GABA, glutamate and acetylcholine (*75–77*).

### Statistical analysis

Whole-brain and region-level statistical analyses were consistently performed throughout this study. The whole-brain analysis was designed to assess global effects and involved computing spatial Spearman correlations between pairs of parcellated brain maps (e.g., between sample entropy and GABA receptor density). To account for the spatial autocorrelation that brain maps typically exhibit, p-values for these associations were obtained via spin permutation by first projecting parcels onto a spherical surface and then performing 10,000 random rotations with iterative reassignments to generate a null distribution (*78*) .

The region-level analysis aimed to assess local association effects between different brain measures. To this end, linear mixed-effects models were fitted separately at each region by regressing the observed values of a given measure (e.g., sample entropy) onto the observed values of another brain measure (e.g., cortical thickness), with subject information included as a random effect and scan acquisition treated as the unit of measurement. The strength of these associations was summarized using z-statistics alongside their corresponding p-values. In both types of analyses, multiple comparisons were consistently controlled using a false discovery rate (FDR) procedure (*79*).

## Author contributions

Conceptualization: J.R, G.P, GM.D, AL.S. Methodology: J.R, G.P, GM.D, AL.S, D.M. Investigation: J.R, G.P, GM.D, A.S, D.M, M.D, G.A, A.S. Resources:J.P, G.P, G.A. Data curation: J.R, G.P, G.A, AL.S, A.S. Validation: L.X., L.Q., and Y.T. Formal analysis: J.R, G.P. G.A,. Visualization: J.R, GM.D, G.P. Supervision: D.M. G.P. G.A J.R. Writing—original draft: GM.D, G.P, AL.S, J.R. Writing—review and editing: G.A, D.M, A.S, M.D. Funding acquisition: G.P, GM.D. Project administration: G.P, J.R

## Data and materials availability

The data that support the findings of this study are available from the corresponding author, GP, upon reasonable request. Source code and scripts to reproduce the results and figures in this manuscript can be found in: https://github.com/jrasero/meg-excitability-landscape.

